# Migration without interbreeding: evolutionary history of a highly selfing Mediterranean grass inferred from whole genomes

**DOI:** 10.1101/2020.09.03.280842

**Authors:** Christoph Stritt, Elena L. Gimmi, Michele Wyler, Abdelmonaim H. Bakali, Aleksandra Skalska, Robert Hasterok, Luis A. J. Mur, Nicola Pecchioni, Anne C. Roulin

## Abstract

Wild plant populations show extensive genetic subdivision and are far from the ideal of panmixia which permeates population genetic theory. Understanding the spatial and temporal scale of population structure is therefore fundamental for empirical population genetics – and of interest in itself, as it yields insights into the history and biology of a species. In this study we extend the genomic resources for the wild Mediterranean grass *Brachypodium distachyon* to investigate the scale of population structure and its underlying history at whole-genome resolution. 86 accessions were sampled at local and regional scales in Italy and France, which closes a conspicuous gap in the collection for this model organism. The analysis of 196 accessions, spanning the Mediterranean from Spain to Iraq, suggests that the interplay of high selfing and seed dispersal rates has shaped genetic structure in *B. distachyon*. At the continental scale, the evolution in *B. distachyon* is characterized by the independent expansion of three lineages during the Upper Pleistocene. Today, these lineages may occur on the same meadow yet do not interbreed. At the regional scale, dispersal and selfing interact and maintain high genotypic diversity, thus challenging the textbook notion that selfing in finite populations implies reduced diversity. Our study extends the population genomic resources for *B. distachyon* and suggests that an important use of this wild plant model is to investigate how selfing and dispersal, two processes typically studied separately, interact in colonizing plant species.

## Introduction

In population genetic theory, a population is a matter of definition. In the natural world, populations are part of complex geographic mosaics; they can be discrete, continuous, or somewhere in-between; and they can expand and contract, appear and disappear. In short, answering the question “What is a population?” is not straightforward for most species (Waples & Gaggiotti, 2006). Yet answering it, implicitly or explicitly, establishes the link between theory and data and is the first step for many studies in ecology, evolution, and conservation.

The concept of population structure or subdivision subsumes many of the challenges of empirical population genetics (Wright, 1931; Hey & Machado, 2003). Population structure results from the phylogeographic and demographic history of the species (Nieto Feliner, 2014; Charlesworth, 2009), life history traits such as dispersal mode and breeding system (Loveless & Hamrick, 1984), and ecological interactions and adaptation (Linhart & Grant, 1996). Disentangling how these different processes contribute to genetic diversity and differentiation is a major goal of evolutionary biology. As processes are inferred from patterns, for any species the disentangling ideally begins with a thorough analysis of population subdivision, that is, of the spatial and temporal scale of genetic differentiation (e.g. Platt et al., 2010).

In the past years, the resolution at which population structure can be analysed has increased greatly due to the availability of genomic data (Pannell & Fields, 2014). Sampling genealogies along genomes allows to overcome problems due to the high stochasticity of single gene trees (Hey & Machado, 2003). Furthermore, measures of absolute genetic divergence such as d_XY_ can replace or complement more assumption-rich statistics such as F_ST_, which rely on *a priori* definitions of populations and can be confounded by within-population structure or the sampling of close relatives (Putman & Carbone, 2014).

In this study we investigate the scale of population structure in the Mediterranean grass *Brachypodium distachyon* and infer its underlying history of population divergence. *B. distachyon* naturally occurs around the Mediterranean Basin and is a specialist of arid, oligotrophic habitats characterized by recurrent disturbance (San Miguel, 2008; López-Alvarez et al., 2015). It has a fully assembled diploid 272 Mb genome (International Brachypodium Initiative, 2010) and is an established model to study various aspects of grass biology (Vogel, 2016, Scholthof et al., 2018). Its value as a model for plant evolution lies in its Mediterranean distribution, providing a window into an ecologically precious yet understudied region (Thompson, 2005); its belonging to the Poaceae, a plant family dominating terrestrial ecosystems around the globe (Gibson, 2009); and, finally, its genome of intermediate complexity, where numerous gene copy number variants (Gordon et al., 2017) and transposable element polymorphisms (Stritt et al., 2018, 2020) contribute to molecular evolution.

A key question which has emerged from previous population genetic studies with *B. distachyon* concerns the role of seed dispersal in shaping genetic structure. A large-scale microsatellite survey in Turkey reported high genotypic diversity within and little differentiation between populations (Vogel et al., 2009). This was explained through a combination of long-distance seed dispersal and extreme selfing rates, the former shuffling genotypes across the landscape, the latter preventing gene flow between diverged genotypes. Other studies, however, found little diversity within and high differentiation between populations in Turkey (Dell’Acqua et al., 2014) and Spain (Marques et al., 2017), in agreement with the increased genetic drift expected in selfing organisms (Charlesworth, 2003). How-ever, as these studies considered different populations and used different sampling schemes, it remains difficult to tell whether their disagreement reflects different biological realities or study designs.

A second pattern highlighted by previous studies is the sympatry of highly diverged lineages differing in flowering phenology (Tyler et al., 2016; Gordon et al., 2017; Sancho et al., 2018). As in other widespread annual plants (e.g. Engelmann & Purugganan, 2006), genetic variation for the control of flowering is present in *B. distachyon*, in particular in the vernalization pathway preventing the development of flowers in the cold season (Ream et al., 2014). It has been suggested that differences in flowering phenology could explain the lack of admixture between sympatric lineages (Gordon et al., 2017). This hypothesis, however, is difficult to evaluate as long as the role of phenotypic plasticity in flowering and the biogeographic history underlying the large-scale distribution of lineages remain unknown.

Here we extend the genomic resources for *B. distachyon* with whole-genome sequences for 75 Italian and 11 French accessions sampled at local and regional scales. Besides filling a conspicuous geographical gap in the *B. distachyon* collection, our aim was to sample across different spatial scales in order to better understand the scale of population structure in this species. Combining these resources with publicly available data, we describe the scale of population structure in 196 accessions spanning the Mediterranean from Spain to Iraq, infer the underlying history of population divergence, and discuss how the interplay of selfing and dispersal has shaped genetic structure across spatial and temporal scales. Furthermore, we conducted flowering time experiments illustrating the phenotypic plasticity of flowering phenology and suggesting caution when extrapolating from greenhouse experiments to natural populations.

## Material & Methods

### Sampling & Sequencing

719 dry *Brachypodium* plants were collected in 2016 across Italy, in the Languedoc region in Southern France, and in 2018 across the Atlas Mountains of Morocco (Table S1). A representative set of 276 accessions were grown in the greenhouse from single seeds. Their DNA was extracted from leaves with a CTAB-based method (Stein et al., 2001), quality-checked with Nanodrop (NanoDrop Technologies Inc), and quantified with Qubit (Invitrogen).

As *B. distachyon* may be confounded with its morphologically similar sister species *B. stacei* and *B. hybridum* (Catalan et al., 2016), we first identified the species of the 276 accessions using the microsatellite marker ALB165 (Giraldo et al., 2012) and by sequencing the GIGANTEA gene (López-Alvarez et al., 2012) with ABI SOLiD. 143 individuals were identified as *B. distachyon*, 130 as *B. hybridum*, and three as *B. stacei* (Table S1). All 70 accessions collected in Morocco proved to be *B. hybridum* and were not included in this study.

86 out of the 143 genotyped *B. distachyon* accessions were selected for sequencing. In order to reduce the chance of sequencing close relatives, only accessions collected at least 50 meters apart were included. In addition, we also sequenced the reference accession Bd21 as a control and the *B. stacei* accession Cef2 as an outgroup for phylogenetic analyses. Paired-end libraries were prepared with the PCR-free KAPA Hyper Prep Kit (KAPA Biosystems). The 150 bp paired-end reads were sequenced with Illumina HiSeq2500 at the Genomics Facility Basel to a mean coverage of 23 (range 18–38, Table S1). The resulting sequencing data were joined with 57 recently sequenced Turkish accessions (Skalska et al., 2020; mean coverage 36) and with data for 52 accessions originating mainly from Spain and Tur-key (Gordon et al., 2017). To speed up computation and avoid bias due to unequal coverage, the latter reads were downsampled to a mean coverage of 30 using Picard tools v.1.97 (broadinstitute.github.io/ picard). The total number of accessions analysed in this study is thus 86 + Bd21 + 57 + 52 = 196.

### Alignment, variant calling & filtering

Paired-end reads were aligned to version 3.0 of the *B. distachyon* Bd21 reference genome, downloaded from Phytozome 12, with BWA MEM v. 0.7.12-r1039 (Li, 2013). Aligned reads were converted to the bam format and sorted with samtools v. 1.9 (Li et al., 2009), and read duplicates were removed with sambamba (Tarasov et al., 2015). Variants were then called with GATK v. 4.0.2.1. The following hardfiltering thresholds were applied to the combined variants: only SNPs were retained with a quality-by-depth > 20, a Phred score < 20 for Fisher’s exact test for strand bias, and a *Z*-score between −1 and 1 for Wilcoxon’s rank sum test of Alt vs. Ref read mapping qualities. Additionally we removed SNPs in low complexity regions identified with dustmasker (Morgulis et al., 2006), resulting in a total of 8,426,015 hard-filtered SNPs.

### Dealing with artificial heterozygosity due to structural variants

Identifying heterozygous SNPs in *B. distachyon* is complicated by a high number of gene copy number variants (Gordon et al., 2017). If a stretch of DNA is duplicated in a sequenced accession but present as a single copy in the reference, sequencing reads derived from the duplicates are collapsed onto a single region when aligning them to the reference genome. Depending on the divergence of the duplicates, this may give rise to islands of extended heterozygosity (IEH): regions of up to few thousand base pairs with extreme numbers of heterozygous sites (Figure S1). As wrongly identified heterozygosity might severely bias downstream analyses, in particular the estimation of selfing rates, we first quantified the issue by scanning each accession for IEH. An IEH was recorded if there were ten or more heterozygous SNPs along a distance of at least 300 bp. The program implementing this approach (detectIEH.py) is available on github.com/cstritt/popgen.

Because hard filtering proved insufficient to deal with the artificial heterozygosity identified in the previous step, we used a masking approach to obtain high-confidence SNPs, including hetero-zygous sites. For each accession a mask was created indicating the regions in the genome within 1.5 standard deviations of the accession-specific mean read coverage. In this way, an average of 85% of the genome was included per accession (range 76% [Bd1-1] to 88% [Bd21]). In a next step, the individual masks were combined with bedtools (Quinlan & Hall, 2010) to define high-confidence regions as regions with “normal” coverage in at least 90 percent of the accessions. Removing SNPs outside these regions resulted in a set of 5,774,928 SNPs across the 196 accessions. These variants are referred to as high-confidence SNPs in the following.

### Population structure and historical gene flow

Genetic structure was analysed using principal component analysis and ancestry coefficient estimation. For the principal component analysis, the 5,774,928 SNPs in high-confidence regions were used and additionally filtered with the snpgdsLDpruning function of the R package SNPrelate (Zheng et al., 2012). Only SNPs were retained with a minor allele frequency > 0.05 and a composite linkage disequilibrium < 0.4, resulting in 14,485 informative SNPs. The snpgdsPCA function of the same package was used to calculate the components. Ancestry coefficients were estimated with sNMF, as implemented in the R package LEA (Frichot & François, 2015), a method appropriate for high levels of inbreeding as it does not assume Hardy-Weinberg equilibrium. Using the same SNPs as for the PCA, sNMF was run for K values from 2 to 20, with 10 repetitions per K.

A rooted phylogeny was estimated with SNPhylo (Lee et al., 2014). For this analysis we selected synonymous sites in the high-confidence SNP set and additionally filtered them for a minor allele frequency of 0.05, a maximum missing rate of 0.1, and a maximum linkage disequilibrium (r^2^) of 0.4, resulting in 5,432 LD-pruned synonymous sites. The root was set to the *B. stacei* accession Cef2, sequenced for this study. 100 bootstrap replicates were performed to assess the robustness of the tree.

fineSTRUCTURE v. 4.0.1 (Lawson et al., 2012) was used to obtain a more detailed picture of population subdivision in *B. distachyon*. This method leverages haplotype information to infer shared ancestry between individuals. More specifically, it estimates the number and lengths of chunks along the genome of individual *i* coalescing with individual *j* before they coalesce with any other individual. The program output we present is a similarity matrix which indicates the recombination map distance a “recipient” genotype (rows) receives from the “donor” genotypes (columns). As fineSTRUCTURE models linkage disequilibrium and performs best with dense haplotype data, the 8,426,015 hard-filtered SNPs were used after removing heterozygous sites and missing data, resulting in 3,467,964 SNPs. By removing heterozygous sites the accessions are treated as haploid and no phasing is thus required. Recombination distances between loci were obtained from the linkage map of Huo et al. (2011). fineSTRUCTURE was run in its haploid mode, using default settings.

To test whether shared ancestry is better explained by common descent or historical gene flow, TreeMix (Pickrell & Pritchard, 2012) was used to model migration. The high-confidence SNP set was used, with 4,473,162 variable sites after removing missing data. To account for linkage disequilibrium, a block size k of 2,000 SNPs was defined, which corresponds to an average distance of 120 kb in the genome. Models with zero to four migration edges *m* were tested, with 100 independent runs for each *m.* For each model, the run achieving the highest likelihoods was used for model comparison.

### Linkage disequilibrium

Selfing has important consequences for the extent of linkage disequilibrium (LD). The effective recombination rate is 1 – s times slower with selfing because meiotic recombination is mainly reshuffling homozygous alleles (Nordborg 2000), resulting in extensive linkage disequilibrium. On the other hand, population substructure in the absence of gene flow decreases LD as mutations accumulate in independently evolving lineages (Hartfield et al., 2018). We estimated linkage disequilibrium at different scales of population structure with PopLDdecay (Zhang et al. 2019). LD is reported as r^2^ averaged over pairwise comparisons. The high-confidence SNP set was used with default program settings (-MAF 0.05, -Miss 0.25). To assess the effect of population subdivision on LD, we used the sum of terminal branch lengths in subtrees as a measure of how much independent evolution has occurred in different subtrees.

### Estimation of divergence times

To put a timescale on the divergence in *B. distachyon*, we used the multispecies coalescent approach implemented in BPP v.4.2.9 (Rannala & Yang, 2003; Flouri et al., 2018). 20 diverse accessions were selected to represent the *B. distachyon* phylogeny. 200 random genomic regions of 1 kb length and at least 100 kb apart were chosen, as BPP assumes no recombination within and free recombination between loci. For each of the 20 accessions the 200 sequences were obtained by calling consensus sequences from the respective bam file. Potential problems of paralogy were avoided by considering only reads with a mapping quality > 30, which excludes reads mapping to multiple places in the genome. Inverse gamma priors were set to (3, 0.014) for the root age τ and to (3, 0.002) for the population size parameter θ, which corresponds to a mean theta of 0.001 and a mean root age of one million years, assuming a constant mutation rate of 7×10^−9^ substitutions per site per generation (Ossowski et al. 2010). The rooted species tree was defined as (((A_East, A_Italia), (B_East, B_West)), C_Italia), as inferred with the SNPhylo analysis described above. The MCMC was run four times independently, each time with 408,000 iterations, including a burn-in of 8000 iterations and samples taken every second iteration. Convergence of the four runs was assessed with Tracer (Rambaut et al., 2018).

### Genetic diversity and population differentiation

To estimate genetic diversity and differentiation, we estimated absolute genetic differentiation d_XY_ at synonymous sites. The --includeNonVariantSites option of GATK v.3.4 was used to obtain variable and non-variable synonymous sites. For each pairwise comparison between the 196 accessions, we then calculated the number of variable and the total number of sites. Heterozygous sites were ignored and only sites were considered for which both accessions in the comparison had a read coverage within 1.5 standard deviations of the accession-specific mean coverage. The script implementing this procedure (pairwise_nucdiff.py) is provided on github.com/cstritt/popgen. Pairwise geographic distances were calculated as Great Circle (WGS84 ellipsoid) distances between each pair, using the spDistsN1 function of the R package sp (Pebesma & Bivand, 2005). Mantel tests were used to test for isolation by distance, using the mantel.rtest function of the ade4 package with 100 permutations to assess the significance of the correlations (Dray & Dufour, 2007). To test if genetic diversity differs between regions, a Kruskal-Wallis test was performed followed by Dunn’s test for multiple comparisons with a Benjamini-Hochberg correction.

### Flowering time experiments

Two experiments were conducted to investigate how flowering phenology relates to genetic structure in *B. distachyon.* Firstly, we tested how the newly sampled accessions respond to vernalization treatments ranging from 0 to 12 weeks at 4°C under short-day conditions in the greenhouse. Twelve Italian acces-sions with different geographic origins were analysed (Cro24, Cm2, Leo2, Mca23, Mca18, Msa27, Ren22, San12, Sap49, Spm16, Sul5, Tar1). We further used four control accessions from Turkey and Iraq (Bd1-1, Bd3-1, BdTR3C, BdTR7a) previously investigated (Ream et al., 2014; Sharma et al., 2017) in order to confirm the reproducibility of the flowering behaviour and vernalization requirements. Plants from all 16 accessions were germinated for three weeks in the greenhouse. They were subsequently put in a cooling chamber for 0, 4, 6, 8, 10 or 12 weeks, respectively, and transferred to the greenhouse again. Three to five replicates were used per accession per time point. Time until flowering was counted from the day after the return to warm conditions (directly after the first three weeks of germination for the plants with 0 weeks vernalization). Flowering was defined as the emergence of the first spike. The cooling chamber was set to 4°C, 8 h light and 16 h dark. The greenhouse conditions were 16 h day at 20–21°C and 8 h dark at 18°C with a light intensity of 200 umol PAR and 60% relative humidity over the entire experiment.

In the wild, plants all go through a prolonged vernalization process during winter which might produce different outcomes than the conditions tested in the greenhouse. To test for this possibility, we planted seeds of 105 accessions outdoors in November 2017, in the Botanical Garden of Zurich, Switzerland, with six replicates per accession. Only six plants did not germinate and 624 plants were used for phenotyping. To ensure no bias due to position effect, replicates were distributed randomly across trays. Mild heating was applied at the bottom of the compartment to prevent freezing when outdoor temperature fell below –10° C. From April 2018 on, plants were checked every three to four days for flowering. The first plant flowered the 21st of April 2018, which was used as a reference (day 1). About 14% of the replicates from the A lineage desiccated before the end of the experiment and did not reach flowering. To test for differences in flowering time between clades a linear mixed effects model was used, as implemented in the R package MCMCglmm (Hadfield, 2010). To control for the six biological replicates, individual accessions were treated as random effects in the model.

## Results

### Occurrence of *Brachypodium distachyon*

*B. distachyon*, *B. stacei*, and their allotetraploid hybrid *B. hybridum* are now known to be different species (Catalán et al., 2012), while they were considered different cytotypes of *B. distachyon* until 2012. As botanical collections do not distinguish between the three species, their relative distribution around the Mediterranean and beyond is still vague. We found that, in agreement with general habitat descriptions (Catalán et al., 2016), *B. hybridum* was more frequent in the drier South of Italy, while *B. distachyon* was predominant in Northern Italy and France (Table S1). Only three plants were genotyped as *B. stacei*, in agreement with this species being the rarest of the three (Table S1). Furthermore, we found only *B. hybridum* in Morocco, suggesting that *B. distachyon* might be rare on the African continent.

### Single nucleotide polymorphisms: beware of artificial heterozygosity

As mentioned above, a significant proportion of the initial 8,426,015 hard-filtered SNPs were observed to aggregate in islands of extended heterozygosity (IEH, Figure S1). IEH can be artefacts which arise when reads from paralogous stretches of DNA are collapsed onto a single region in the reference genome; but they can also represent real heterozygosity in the case of recent admixture between diverged lineages. To distinguish the two possibilities, we compared IEH among the initial hard-filtered SNPs and 5,774,928 SNPs in high-confidence regions, defined as regions where at least 90 percent of the accessions have a read coverage within 1.5 standard deviations of their mean coverage. If a region has a coverage within this “normal” range, this indicates that it was not deleted or duplicated in the sequenced accession, and that heterozygous sites are thus not due to collapsed paralogs.

A median of 116 IEH were identified per accession in the hard-filtered set as compared to only two in the high-confidence set (Figure 1), indicating that most IEH are artefacts caused by structural variants. Six accessions (Uni2, Mig3, ABR9, Bd21-3, ABR7, Per1) deviate from the general pattern and have hundreds of IEH even in high confidence regions (Figure 1a). As the sequencing depth at these IEH is within the normal range and does not suggest the presence of structural variation (Figure S2), we propose that this is real heterozygosity left over from recent interbreeding between diverged genotypes. On average 46% of the IEH identified in any accession overlap with annotated genes, with a median of 50 genes per accession showing evidence of extended heterozygosity. As expected from the composition of the pan-genome (Gordon et al., 2017), the total set of 4,211 genes with IEH is enriched for the GO terms “defence response”, “response to stress” and other functions associated with highly dynamic gene families.

**Figure 1.**
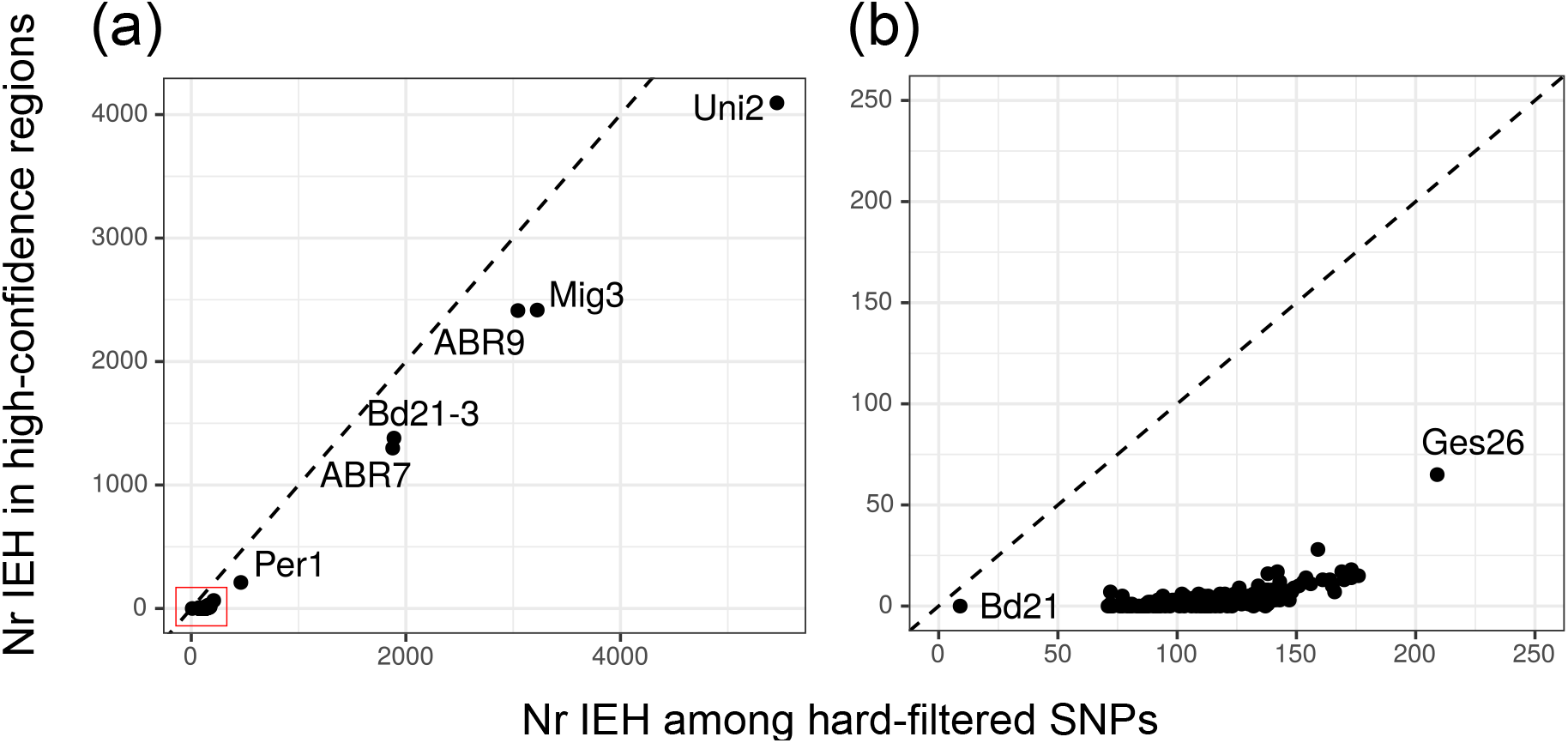
Islands of extended heterozygosity (IEH) among hard-filtered SNPs (x axis) and among SNPs in high confidence regions (y axis), defined as regions where at least 90 percent of the accessions have a read coverage within 1.5 standard deviations of their mean coverage. (a) Overview of all 196 accessions, highlighting the six outlier accessions. (b) Zoom-in on the red square in panel a.

### Three major lineage and five large geographic clades of *B. distachyon*

The different clustering methods capture a clear subdivision of the 196 accessions into five major genetic clades, except for two accessions from the Pyrenees which cannot be assigned (Arn1 and Mon3; Figure 2a). This subdivision is particularly clear in the sNMF analysis, where model fit improves in large steps until K=5, after which it declines in smaller steps as regional and local populations are recovered (Figure 2b), as also shown in an Evanno-like evaluation of the “best” K (Figure S3; Evanno et al., 2005). At the base of the rooted *B. distachyon* phylogeny (Figure 2c,e; Figure S4; an interactive tree/map is provided on github.com/cstritt/bdis-phylogeo), 15 Italian accessions form a clade (hereafter called C_Italia) consisting of nine plants from the Gargano peninsula in south-eastern Italy and six col-lected across Italy, namely in Sicily, southern Campania and Liguria (Figure 2a). After the divergence of this basal lineage, the tree splits into two large lineages: one comprising all the remaining plants from Italy (A_Italia) and plants primarily from the coastal regions of Turkey (A_East), the other comprising plants from the western Mediterranean (B_West) and plants primarily from the Anatolian plateau and south-eastern Turkey/Iraq (B_East). The same subdivision is evident in the PCA (Figure 2d). A geographic outlier are three accessions from southern Turkey (2_20_16, 2_14_15 and 2_14_20; Figure 2a) which cluster with accessions from Spain and France. As these Turkish accessions were collected recently (Skalska et al., 2020) and not grown in contact with Spanish or French accessions, this finding can hardly be explained by mislabelling or seed contamination.

**Figure 2.**
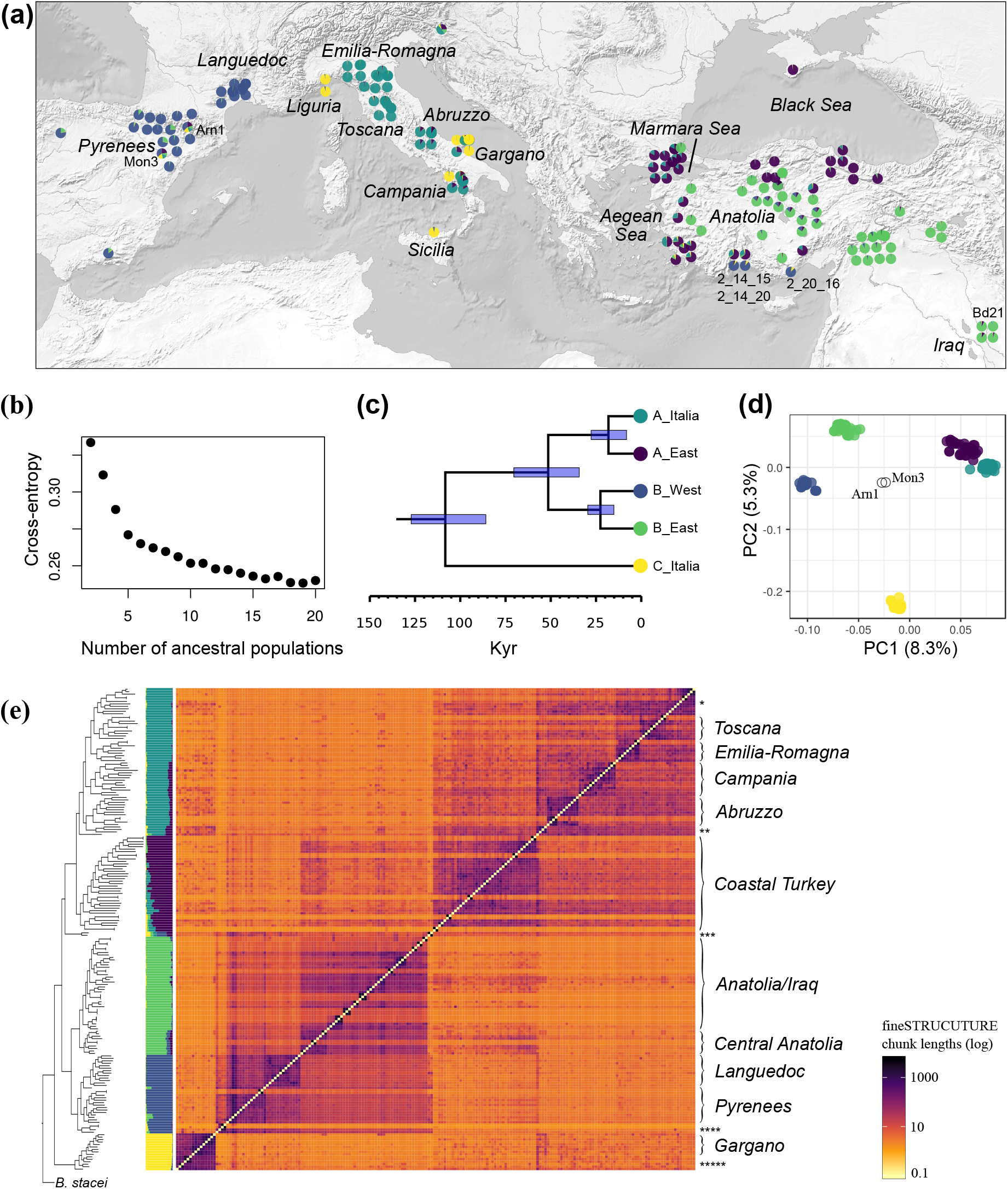
Geographic origin and genetic structure. (a) Map and pie charts showing shared ancestry. Only a subset of accessions is displayed for the dense local samples in Italy and France. (b) Cross-entropy for increasing K in the ancestry analysis with sNMF. (c) Rooted phylogeny with divergence times, estimated under a multispecies coalescent model. (d) Principal component analysis. (e) Rooted phylogeny and coancestry matrix estimated with fineSTRUCTURE, as well as sNMF barplot for K=5. The fineSTRUCTURE chunk length is the recombination map distance donated to individuals in rows from individuals in columns (log scale). Names in italics highlight regional populations, asterisks indicate small clusters: * Accessions from northern Italy, including a cluster of five accessions from Taro valley in Emilia-Romagna. ** ABR9 collected in Albania and three accessions from the Gargano peninsula. *** Arn1 and Mon3. **** Three Turkish accessions clustering with Bd30-1 from southern Spain. ***** Accessions from Sicily, Campania and Liguria. An interactive phylogenetic tree/map can be explored on github.com/cstritt/bdis-phylogeo.

Using the multispecies coalescent approach implemented in BPP (Figure 2c), the split between the basal C lineage and the others is dated to 107 thousand years ago (kya, 95% highest posterior density interval [HPDI] 86–128 ky) and the split between the A and B lineages to 52 kya (95% HPDI 35–72 ky). Within the A lineage, the separation between A_East and A_Italia is dated to 18 kya (95% HPDI 8– 27 ky), while within the B lineage the separation between B_East and B_West is estimated at 22 kya (95% HPDI 15–30 ky).

### Little evidence for historical gene flow between diverged lineages

At this coarse level of genetic structure, we thus observe an incongruence between genetic structure and geographic occurrence. Roughly half of the plants collected in Turkey and Iraq are more closely related to plants from the western Mediterranean than to other plants from the east. Similarly, in Italy the majority of the collected plants are more closely related to plants from the east than to the 15 Italian plants at the base of the phylogeny. In the east, the A and the B lineage might occur in the same locality, yet there is a clear overall geographic division (Figure 2a; github.com/cstritt/bdis-phylogeo; see also Skalska et al., 2020): lineage A is predominant in the coastal region, lineage B on the inland Anatolian plateau and in the Fertile Crescent. In Italy the occurrence of the A and the C lineage is less clear-cut: in two of the four localities where lineage C was found, plants of the A and the C lineage were present on the same meadow and thus sympatric in a narrow sense (Figure S5).

Estimates of shared ancestry show little evidence for recent gene flow between co-occurring lineages in Turkey and Italy (Figure 2a,e). Historical gene flow between geographic clades appears possible, yet the evidence is not particularly strong: a tree-like model without migration edges explains 99.7% of the covariance of allele frequencies between geographic clades (Figure S6), suggesting that shared ancestry apparent in Figure 2 is due to common descent and shared genetic drift rather than admixture. Adding a migration edge leading from A_East to B_East leads to an increase to 99.98%, suggesting that, if historical gene flow has occurred, it has not left strong signatures.

### High selfing rates and a strong effect of subdivision on linkage disequilibrium

Selfing rates often vary within species (Barrett, 2003; Griffin & Willi, 2014), yet studies with *B. distachyon* have all reported a near absence of heterozygosity (Vogel et al., 2009; Marques et al., 2017). To test whether this pattern holds across geographic regions, we estimated inbreeding coefficients and self-ing rates for the 196 accessions. Using SNPs in high-confidence regions, as defined above, we estimated a median inbreeding coefficient F of 0.996 among the 196 accessions (mad=0.003, Table S1), which translates to a median selfing rate of 0.998 using s = 2F / 1+F. When population structure is taken into account and F is estimated within geographic clades, median F is 0.991 and thus only marginally lower than at the species level. As expected, lower F values are estimated for the six accessions whose genomes contain numerous IEH. Overall, our results show that extreme levels of homozygosity and high selfing rates predominate throughout the species’ range and vary little between subpopulations.

LD is low at the species level (Figure 3a) and successively increases as one considers finer scales of subdivision (Figure 3b–d). A comparison of LD decay and the topology of the *B. distachyon* phylogeny supports the expectation that LD decay reflects substructure rather than recombination. Using the sum of terminal branch lengths in subtrees as a measure of how much independent accumulation of mutations has occurred at different levels, we observe a strong negative relation between mean r^2^ at a distance of 100 kb and the sum of terminal branch lengths (Figure S7): LD decay is slower in subtrees containing either few leaves, e.g. in the C lineage, or containing short terminal branches, as in B_East, where there is a surprising number of genetically highly similar accessions.

**Figure 3.**
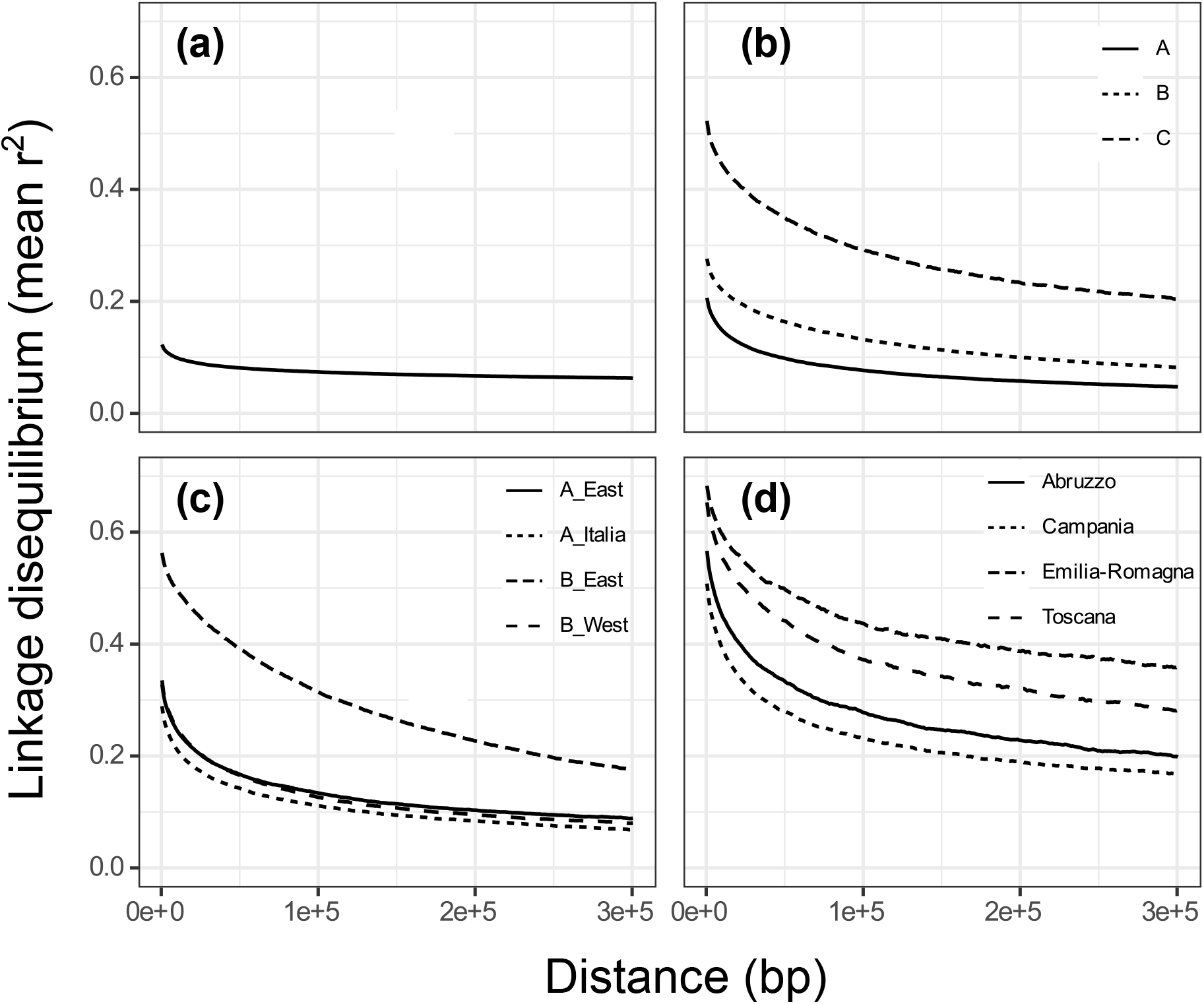
Decay of linkage disequilibrium at different scales of population structure: (a) species-wide, (b) in the three major genetic lineages, (c) in four geographic clades (C_Italia is not included as it is the sole representative of lineage C and thus already included at the lineage level), (d) four regional populations in Italy.

### Strong isolation by distance within the three major lineages

To understand how genetic diversity is partitioned across space and the different levels of genetic structure in *B. distachyon*, we estimated pairwise genetic differences (d_XY_) at synonymous sites. Within the five geographic clades, median d_XY_ is 0.0018, that is, 1.8 different sites per 1000 bp (Figure 4a). Values differ somewhat among clades: d_XY_ is higher in A_East (0.0019) than in the other lineages (P<2×10^−16^, Kruskal-Wallis), and lower in C_Italia (0.0017) than in the others (P<0.03). Levels of diversity in the B_East and B_West clades are similar (0.0017, P=0.37). Between clades, d_XY_ reaches up to 0.0065 in comparisons between C_Italia and B_West.

**Figure 4.**
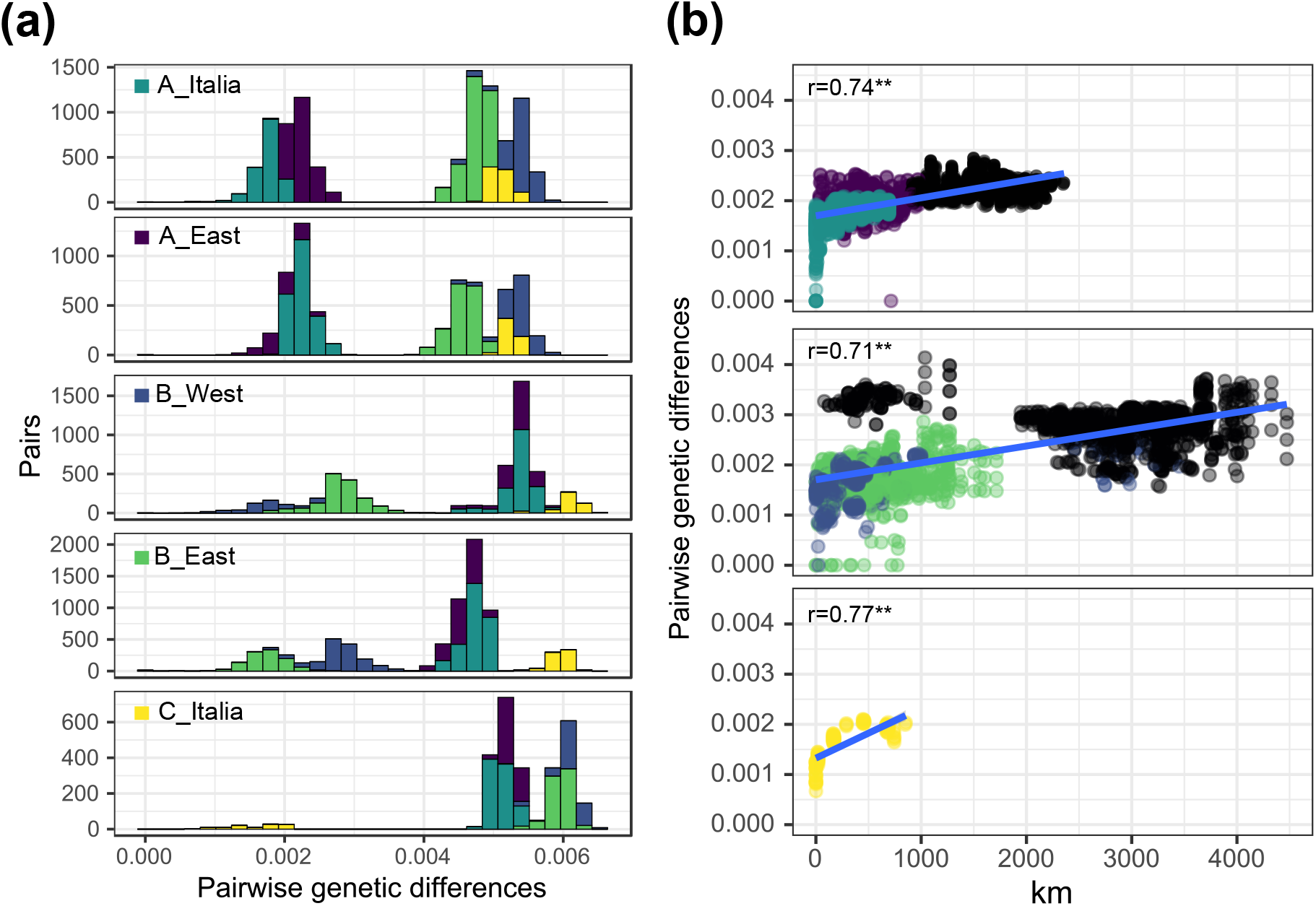
Genetic diversity and differentiation. (a) Pairwise genetic differences (dXY) within and between the five geographic clades. One panel is shown for each clade, the colours within the panel showing with which clade pairwise comparisons are made. (b) Correlation between geographic and genetic distance within the three major genetic lineages A, B and C. Colours within the panels indicate between which geographic regions pairwise comparisons are made. Correlation coefficients from Mantel test and their significance are displayed.

Within all three major lineages, d_XY_ increases with geographic distance, as shown by significant Mantel correlations (Figure 4b). While correlation coefficients are similar between lineages (r_A_=0.75, r_B_=0.71, r_C_=0.77), some deviations are observed in the B lineage. On the one hand, there are genetically similar accessions with distant geographic origins, particularly in the B_East clade, in agreement with the previously observed poor mapping between geography and phylogeny in this clade. On the other hand, genetically distant individuals can be geographically close (upper left of the middle panel in Figure 4b). These are comparisons of Turkish accessions with the three accessions collected in Turkey clustering genetically with B_West accessions (Figure 2a).

### The hierarchical nature of population subdivision

Population subdivision in *B. distachyon* is hierarchical and does not end at level of the five geographic clades. Substructure is evident for increasing values of K in the sNMF analysis (Figure S8), in within-lineage PCAs (Figure S9), as subtrees in the phylogeny (Figure S4), and most clearly in the fineSTRUCTURE analysis (Figure 2e). The clustering algorithm implemented in fineSTRUCTURE identi-fies 59 populations with a median size of two (range 1-13). This inflated number of populations most likely reflects the limitations of formal approaches to identify the “most realistic” number of populations in a highly selfing organism, where the homogenizing force of gene flow is limited and populations are better understood as phylogenetic clades than as panmictic units.

In *B. distachyon*, many subtrees cluster geographically into regional populations (Figure 2e; github.com/cstritt/bdis-phylogeo). In the A_Italia clade, populations from Campania, Abruzzo and Gargano in the South and from Toscana and Emilia-Romagna in the North can be distinguished. Also in the other lineage present in Italy, C_Italia, the 15 samples cluster according to their geographic origin. In the western Mediterranean, the Pyrenees separate a Spanish from a French population comprising all the newly collected accessions from the Languedoc region. Substructure is less obvious in the east. A_East is a diffuse clade occupying primarily coastal areas in Turkey. In the B_East clade, ten accessions from central Anatolia form a well-supported group at the base of the clade. The rest of the samples cover a large area from Baghdad to the Marmara Sea region and do not show an obvious mapping between phylogeny and geography.

### Local and regional patterns of genetic diversity in Italy

The local and regional sampling in Italy allows us to take a closer look at how genetic diversity is distributed across small spatial scales. In Italy, the Apennines separate *B. distachyon* into genetically differentiated regional populations (Figure 5a). We here focus on four regional populations belonging to the A_Italia clade for which the largest number of samples is available (7-14). A TreeMix analysis with these populations suggests that no gene flow has occurred across the Appenines, as a tree-like model without migration edges explains 99.99% of the covariance of allele frequencies between populations (Figure S10).

**Figure 5.**
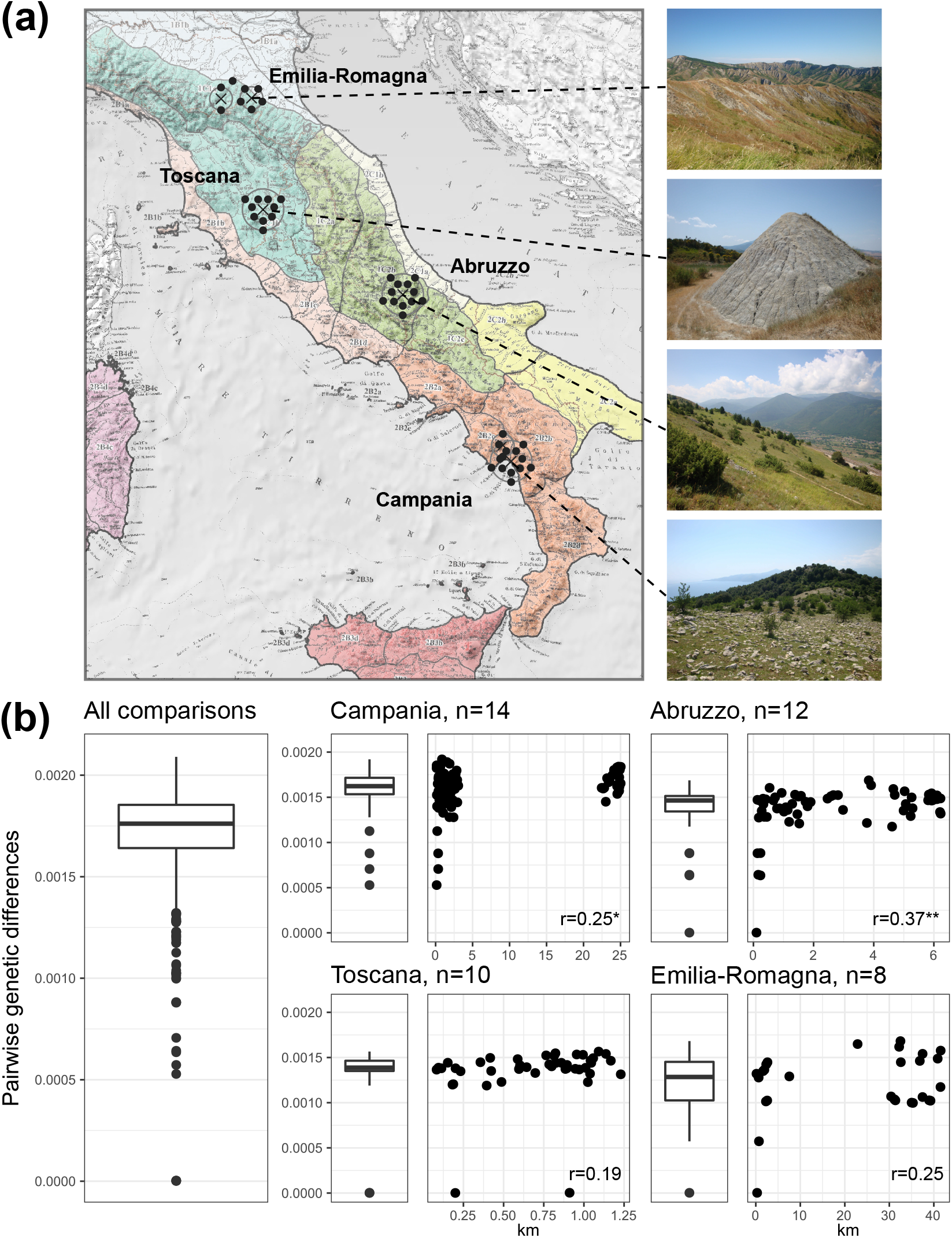
Genetic differentiation across ecoregions in Italy. (a) Map showing the ecoregion classification of Blasi et al. (2014) and *B. distachyon* habitats in the four sampled regional populations. (b) Genetic diversity within the A_Italia clade, for which the most extensive sampling is available. Boxplots show the distribution of d_XY_ across the whole clade and within regional populations, while dotplots show how d_XY_ relates to geographic distance.

Figure 5b shows pairwise genetic differences in the four populations, for which samples were collected few dozen meters to few kilometres apart. Within populations, some more closely related individuals stand out (d_XY_ < 0.001); these were typically collected within 1 km of each other. Most plants, however, even if growing close-by, are genetically quite distinct and can be more different from each other than from accessions kilometres away. Isolation by distance is weak at this scale and, while still significant in the two southern populations, is not significant in the Toscana and Emilia-Romagna populations. Genetic diversity declines step-wise from south to north, with a median d_XY_ of 0.0016 in Campania, 0.0015 in Abruzzo, 0.0014 in Toscana, and 0.0013 in Emilia-Romagna (Figure 5b).

### Flowering phenology in the greenhouse and outdoors

Previous studies described the co-occurrence of the A_East and B_East clades in Turkey and interpreted the lack of gene flow between these lineages as a result of variation in flowering time: acces-sions from the A_East clade require long vernalization and show a delay in flowering under greenhouse conditions (Tyler et al., 2016; Gordon et al., 2017).

Our greenhouse experiment replicated the previously identified flowering behaviour of the two B_East accessions Bd3-1 and BdTR3C and the two A_East accessions Bd1-1 and BdTR7a, which in this order required four, six, eight, and more than 12 weeks to satisfy the vernalization response (Figure S11a, Table S2). With the exception of two C_Italia accessions which required six weeks of cold, all other 10 newly tested Italian accessions required long periods (>10 weeks) of cold followed by up to 40 days in warm conditions to flower (Figure S11a, Table S2).

Flowering behaviour in the outdoor experiment was quite different (Figure S11b, Table S2). All plants emerged in autumn and flowered next spring within two weeks. However, clades still differ in their flowering behaviour. C_Italia and B_East accessions tended to produce flowers more rapidly than A_East accessions, though this difference is not statistically significant at the 5% level. A_Italia and B_West, on the other hand, take longer to flower than the other clades (LME, P< 0.006 for all comparisons between these two clades and the others), even though overall only two weeks separate the early from late flowering accessions.

## Discussion

*B. distachyon* was initially developed as a model to study gene function in grasses, with an emphasis on applications in crop breeding. With this study we extend the genomic resources for this Mediterranean grass and lay a foundation for evolutionary studies by providing a historical explanation for genetic structure in the species. Population structure in *B. distachyon* is strongly hierarchical. It originated during the Upper Pleistocene (c. 129,000 −11,700 years ago) and reflects, as we will argue in the following, the combined effect of high selfing rates and effective seed dispersal. At the continental scale, dispersal and selfing help to explain why highly diverged lineages might be present in the same locality without signs of admixture. At the local scale, selfing and dispersal interact and maintain high local genetic diversity, with important implications for how natural selection is expected to act on the species.

### Extreme selfing and extended heterozygosity as a result of recent outcrossing

In plants, the wide occurrence of sexual reproduction and the elaborate means evolved to ensure it suggest that self-fertilization is disadvantageous under a wide range of conditions (Barrett, 2010). The best known exception are colonizing plants (Baker, 1955; Stebbins, 1957): here, the advantage of reproductive assurance during colonization can outweigh the costs of inbreeding. *B. distachyon* clearly is a colonizing plant: it thrives in disturbed areas like pastures or along paths and declines when stable conditions allow the establishment of perennial competitors (San Miguel, 2008).

Considering this life style, the highly reduced cleistogamous flowers (Khan & Stace, 1999; Dinh Thi et al., 2016), and the general pattern that wind-pollinated species tend to be either predominantly outcrossing or selfing (Barrett & Eckert, 1990), the highly selfing nature of *B. distachyon* comes at no surprise. Nevertheless, the degree of homozygosity revealed by molecular markers is astonishing. In a large-scale survey of 187 accessions from Turkey with 43 microsatellite markers, only four heterozygous sites in four individuals were detected (Vogel et al., 2009). Similarly, Marques et al. (2017), looking at 137 plants in 14 populations with 10 markers, estimated inbreeding coefficients of 1 in all except one population.

Our study agrees with this finding and suggests that high selfing rates are prevalent throughout the species’ range. It also corrects previous estimates of heterozygosity based on genome-wide SNPs, where we reported a median of 7.8 and 6.3% of heterozygous SNPs in the B_East and B_West clades (Bourgeois et al., 2018). These estimates were most likely too high as we did not take into account artificial heterozygosity due to structural variation. Indeed, distinguishing real from artificial heterozygosity proved to be challenging. As we show here, a method based on the clustering of heterozygous sites and sequencing depth can be used to identify artificial heterozygosity arising from deletions and duplications. As plant population genomics is extending to more complex and dynamic genomes, searching for islands of extended heterozygosity can be a simple first step to identify problems with heterozygous sites and to limit biases in downstream analyses.

While self-fertilization appears to be the norm in *B. distachyon*, we also find evidence for recent outcrossing. A large part of the heterozygosity we observed occurred in only six accessions, in the form of extended islands of heterozygosity. Extended heterozygosity can result from the purging of deleterious homozygosity, as observed in inbred maize (Brandenburg et al., 2017), or from balancing selection (Nordborg et al., 1996). These processes, however, imply a selective advantage of heterozygosity and would hardly create hundreds of IEH in single accessions as observed here, but rather few IEH present at higher frequencies. Our results thus provide evidence that the IEH in the six accession are due to the admixture of diverged genotypes. As heterozygosity disappears within a few generations of selfing (Hedrick, 2011), the outcrossing events leading to the six heterozygous accessions must be recent and might have occurred during propagation of the accessions for research and seed distribution.

### Shared ancestry reflects common descent rather than admixture

Historical gene flow can be inferred when the covariance of allele frequencies between populations is poorly explained by tree-like evolution and genetic drift (Pickrell & Pritchard, 2012). Our tests for gene flow between the five geographic clades and between regional populations in Italy suggest that patterns of shared ancestry can largely be explained by tree-like descent rather than historical admixture. These results contrast with previous studies which have interpreted shared ancestry as admixture (Gordon et al., 2017; Stritt et al., 2018) and inferred frequent introgressions from incongruences between plastome and nuclear phylogenies (Sancho et al., 2018). In the latter study, several B_East accessions are shown to have B_West plastome genotypes, but no admixture is evident at the nuclear level, which the authors explain through repeated backcrossing and loss of the B_West nuclear ancestry. Here too, however, common descent and incomplete lineage sorting of plastome haplotypes are a plausible alternative explanation, as for example described for *Hordeum* (Jakob & Blattner, 2006).

While outcrossing and historical gene flow remain a possibility, our results suggest that alternative explanations for shared ancestry, more compatible with the extreme homozygosity and high selfing rates observed throughout *B. distachyon*, should be considered in future studies.

### A scenario of separate migration corridors

To our current knowledge, *B. distachyon* in the Mediterranean region comprises three highly diverged genetic lineages which can be further divided into geographic clades and regional populations. This basic tripartition was described previously in a genotyping-by-sequencing study (Tyler et al. 2016), yet a biogeographic scenario to account for the distribution of the three lineages is now only starting to emerge as samples from the entire species range are being collected (Skalska et al., 2020; Gordon et al. 2020). Here we show that the A lineage is present throughout Italy, and we describe a so far poorly known lineage of *B. distachyon*, C_Italia.

The C lineage diverged from the others 86–128 kya and forms the basal branch in the phylogeny. Previous studies included single samples of this lineage, all originating from Sicily (Tyler et al. 2016, Gordon et al. 2020). We found this lineage to be present yet rare throughout Italy, in two instances growing on the very same meadow as plants belonging to lineage A. Recently a similar situ-ation has been described for *B. hybridum*, where individuals with a common ancestor more than one million years ago coexist (Gordon et al., 2020; Shiposha et al., 2020). *Brachypodium* thus provides striking examples of cryptic grass diversity in the Mediterranean region which would be difficult to appreciate without the help of molecular markers.

The geographic co-occurrence of genetically diverged lineages in *B. distachyon* suggests that three lineages of *B. distachyon* have expanded independently in the past, and that today, where their ranges overlap, extreme selfing and possibly other reproductive barriers prevent interbreeding. While the biogeography of the C lineage remains vague because of the low number of samples, an intriguing scenario which could explain the disjunct genetic structure in *B. distachyon* is that lineages A and B used different migration corridors: a North African corridor for lineage B, and a European corridor for lineage A. To our knowledge, all samples collected in the Balkans so far belong to the A or the C lineage (Tyler et al., 2016; Gordon et al., 2020; our unpublished data), suggesting that the B lineage is absent or rare between the Ligurian Alps and inland Turkey. With additional samples from the Balkans, the Middle East, southern Spain and Africa, some of which have already been sequenced, this scenario should be testable in the near future.

Our molecular dating analysis and the strong isolation by distance observed at the lineage level suggest that the three lineage expansions in *B. distachyon* are not recent and might predate the end of the Last Glacial Period c.11,700 years ago. The timescale we infer in this study, with a time to the most recent common ancestor of 86 to 128 years ago, hinges on an assumed mutation rate of 7×10^−9^, obtained from an experimental evolution study with *A. thaliana* (Ossowski et al., 2010). This estimate is at the fast end of the spectrum as it reflects the accumulation of mutations before purifying selection had a chance to remove them (Ho et al., 2011). Our estimates are therefore more likely too recent than too old. A similar estimate, 162 ky to the common ancestor of the three lineages of *B. distachyon*, was recently obtained by time-calibrating an internal node rather than assuming a mutation rate (Gordon et al. 2020), providing further support for the timescale inferred here.

### Genetic structure and flowering time differences

Previous studies have demoted geography to “a secondary factor in shaping population structure” (Gordon et al., 2017) and instead identified flowering time differences as “a main factor driving rapid intraspecific divergence” (Sancho et al., 2018; see also Tyler et al., 2016). This view is problematic as it is based on flowering phenotypes observed in the greenhouse. Flowering is a plastic trait and depends on environmental conditions (Gaudinier & Blackman 2020) and the timing of the other major life history event in annual plants, germination (Martínez-Berdeja et al., 2020). In our outdoors experiment, all accessions produced flowers within two weeks when they went through prolonged vernalization during winter. While this experiment does not reflect all the environmental variation present in *B. distachyon* natural habitats, it suggests that differences observed in the greenhouse might be exaggerated through artificially short vernalization times.

While the role of flowering in reproductive isolation is less clear than suggested by previous studies, flowering remains an intriguing trait in *B. distachyon*. A geographic pattern of flowering time differences is observed in Turkey but not in Italy. In Turkey, the A lineage, which requires vernalization to flower, is present along the coast, while the B lineage, which requires little to no vernalization (Gordon et al. 2017, Skalska et al. 2020, this study), is mainly found in inland Anatolia (Figure 2). Skalska et al. (2020) recently showed that the two lineages also differ in drought tolerance, indicating that the B_East accessions are adapted to the arid climate of central Anatolia. In Italy, no such geographic differences in vernalization requirement and flowering are observed, neither among lineages nor among regional populations. Possibly this reflects the stronger ecological contrasts in Turkey, where rain is scarce on the inland Anatolian plateau even in winter, while the coastal regions have the typical Mediterranean winter rainfalls (Sensoy, 2008). Investigating the molecular evolution of flowering genes in *B. distachyon* will allow to further characterize the putative adaptive role of flowering time in this species.

### Local consequences of selfing and dispersal

Within regional populations in Italy, isolation by distance is weak, and sampling within few hundred meters can yield individuals as different from each other as individuals kilometres apart (Figure 5). This pattern is discernible in the distribution of pairwise genetic differences and as long terminal branch lengths in the phylogeny, especially in the A lineage. Clearly the plants collected locally did not descend from single recent founder events, which would result in high local similarity. Our study thus supports the hypothesis that the interplay of high selfing and seed dispersal rates shapes local and regional genetic diversity, as proposed by Vogel et al. (2009) for populations in Turkey.

In population genetics textbooks, migration, gene flow, and interbreeding are often used synonymously (e.g. Charlesworth & Charlesworth, 2012), reflecting the dominance of animal models in the field. In plants, however, migration can occur through pollen and seed, and the migration of seeds need not imply the exchange of genetic material. With a selfing rate close to one, lineages can evolve independently of each other for a considerable period without the homogenizing effect of interbreeding, and lineage-specific mutations may accumulate. Seed dispersal would then ensure that closely related genotypes do not occur in patches, as observed in *A. thaliana* (Bomblies et al., 2010), but are shuffled across the landscape.

Central to the selfing-dispersal hypothesis is that *B. distachyon* disperses its seeds easily and over long distances. Morphological adaptations to do so have been described: the species has barbed palea which facilitates zoochory (Catalán et al., 2016). Surprisingly, however, so far no study mentioned the animals themselves, sheep and goats. From our fieldwork and previous reports (Vogel et al., 2009) it is clear that the habitat in which *B. distachyon* thrives are arid, oligotrophic pastures grazed by sheep and goats. Not only do these animals keep away more competitive perennial plants, but they are also known to carry plant seeds over long distances and to be an important vector of long-distance seed dispersal (Manzano & Malo, 2006). In the Mediterranean landscape, which has been shaped by human activity through thousands of years (Grove & Rackham, 2001), the movement of domestic animals might thus be a crucial factor in the population dynamics of *B. distachyon* and similar plants.

The possibility that genetic diversity is heavily influenced by long-distance seed dispersal has important implications for the study of natural selection in *B. distachyon.* While selfing itself decreases the fixation time of adaptive mutations and might favour local adaptation (Glémin & Ronfort, 2013; Hartfield et al., 2017), selfing combined with high rates of seed migration is expected to favour phenotypic plasticity over local adaptation (Sultan & Spencer, 2002). It is notable that no instance of a single lineage rising to high local frequencies is observed. Instead, it appears that seed dispersal maintains high local genetic diversity and might compensate for what is considered a main disadvantage of selfing: low diversity and failure to adapt to spatially and temporally heterogeneous environments (Wright et al., 2013).

## Conclusion

What is a population of *B. distachyon*? *B. distachyon* is a generalist colonizing grass with large, continuous populations where pastoralism has not given way to intensive agriculture or forest (San Miguel, 2008; observations during fieldwork). As genotypes are shuffled across the landscape, a local sample is likely to capture the genotypic diversity of a broader region. Indeed, such a sample might include highly diverged genotypes coalescing only in the distant past, which might bias analyses relying on a purely spatial definition of “population”. With these properties, *B. distachyon* is an interesting model to explore the limitations of classical population genetic models and to investigate the surprising ways in which the evolutionary play is acted out in nature.

## Acknowledgments

The authors are especially thankful to the Swiss National Science Foundation and the University of Zurich’s Research Priority Program (URPP) *Evolution in Action* for funding this project. AS, LAJM and RH acknowledge the funding from the National Science Centre Poland (grant no. 2015/18/M/ NZ2/00394). The authors would like to thank the Genetic Diversity Center at ETH Zurich for providing access on their High Performance Computing resources as well as Christian Beisel at BSSE-Basel, Gerhard Herren and Helen Zbinden for their support in the lab. The authors also would like to thank Marco Maccaferri, Saverio Sciandrello and Pietro Minissale for their help in sampling in Italy, JeanPaul Peltier for establishing connection between Switzerland and Morocco, Christian Parisod for his comments on an early draft of the manuscript and the three anonymous reviewers for contributing greatly to improving the manuscript.

## Data Accessibility

Illumina paired-end sequencing data generated for this project are available on the European Nucleotide Archive, project number PRJEB40344.

## Author Contributions

ACR, LAJM, RH obtained funding for the project. ACR and CS designed the study. ACR, AHB, CS, NP and RH collected samples. CS analysed the data, with input and help from ACR, AS, ELG and MW. CS wrote the original draft, and all authors reviewed it.

