## Supporting Figures S1-11 for "Migration without interbreeding: evolutionary history of a highly selfing Mediterranean grass inferred from whole genomes"

**Table of contents**


| | |
|---|---|
| Figures S1: Genome browser visualisation of islands of extended heterozygosity | 2 |
| Figure S2: Read coverage in islands of extended heterozygosity | 3 |
| Figure S3: Evanno-like evaluation of the "best" level of population structure | 4 |
| Figure S4: Rooted phylogeny with leaf labels and bootstrap values | 5 |
| Figure S5: Sympatry of diverged lineages | 6 |
| Figure S6: TreeMix analysis to test for gene flow between the five geographic clades | 7 |
| Figure S7: Relation between LD decay and terminal branch lengths in subtrees | 8 |
| Figure S8: sNMF analysis with K from 2 to 20 | 9 |
| Figure S9: Within-clade PCAs | 10 |
| Figure S10: TreeMix analysis to test for gene flow between regional populations in Italy | 11 |
| Figure S11: Flowering phenology in the greenhouse and under semi-natural conditions | 12 |


Table S1, containing information about the samples (coordinates, genotyping, sequencing statistics, selfing rates), is provided as a separate file, as well as table S2 containing the tabulated results of the two flowering time experiments.

**Figure S1:** Genome browser visualisation of islands of extended heterozygosity. Plots show paired-end reads aligned against the reference genome of two accessions: the reference itself (Bd21) and the two accessions BdTR7a and Uni2. a) An IEH caused by a gene copy number variant. Heterozygosity coincides with a doubling of sequencing depth. b) An IEH showing real heterozygosity due to recent admixture. Here heterozygosity stretches over long distances and the sequencing depth is within the normal range.

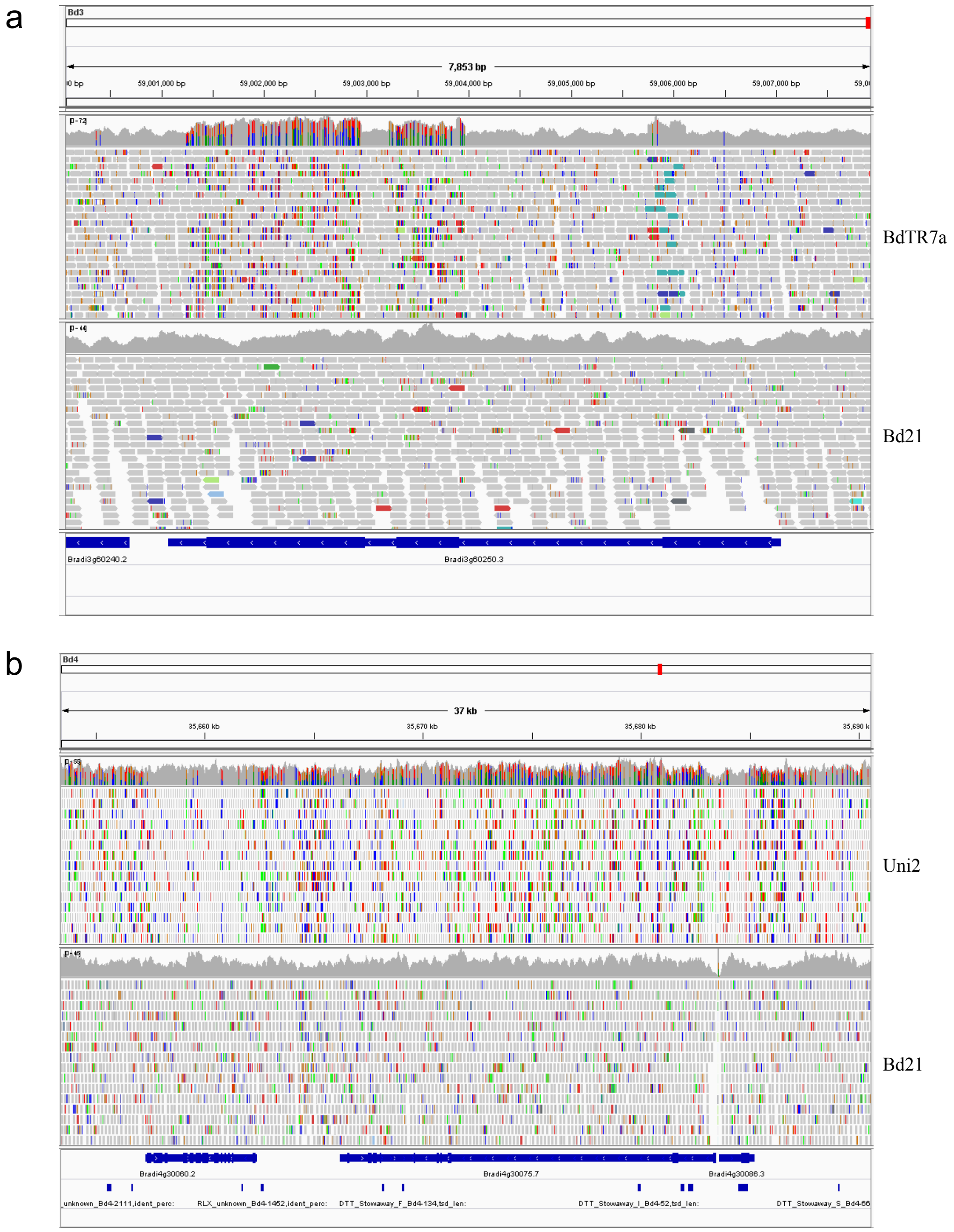

**Figure S2:** Read coverage in islands of extended heterozygosity in a) the six outlier accessions and b) ten random accessions. Red crosses show the median read coverage across the genome for each accession, while the boxplots show read coverages in IEH.

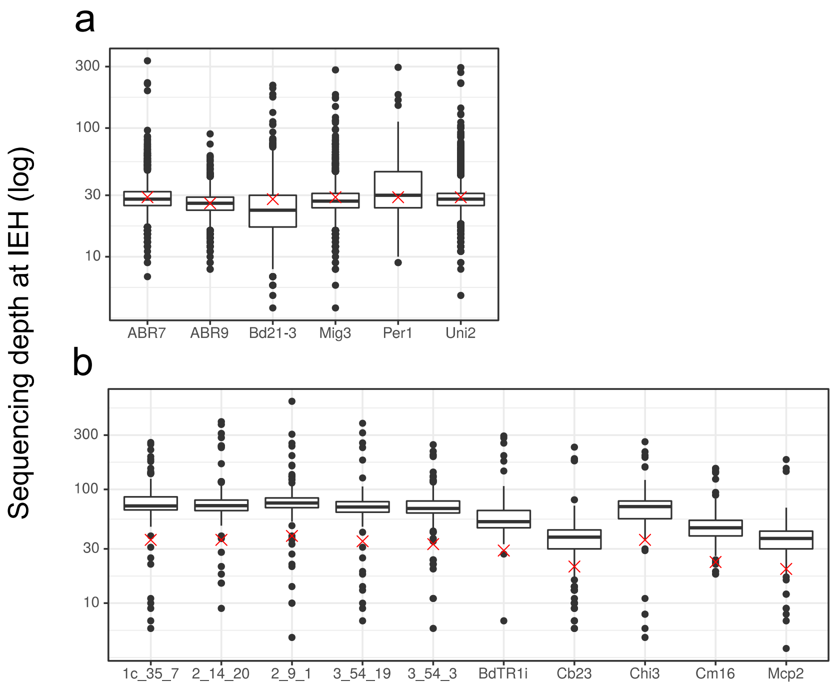

**Figure S3:** Evanno-like evaluation of the most conspicuous level of population structure. The same calculations are performed as described in Evanno et al. 2005, with the difference that likelihoods are replaced by cross entropy (CE), the measure of model fit used by sNMF. Distributions for single values of K represent the 10 repetitions performed for each K. As discussed in the article, high selfing rates suggest caution in the interpretation of the "best" K.

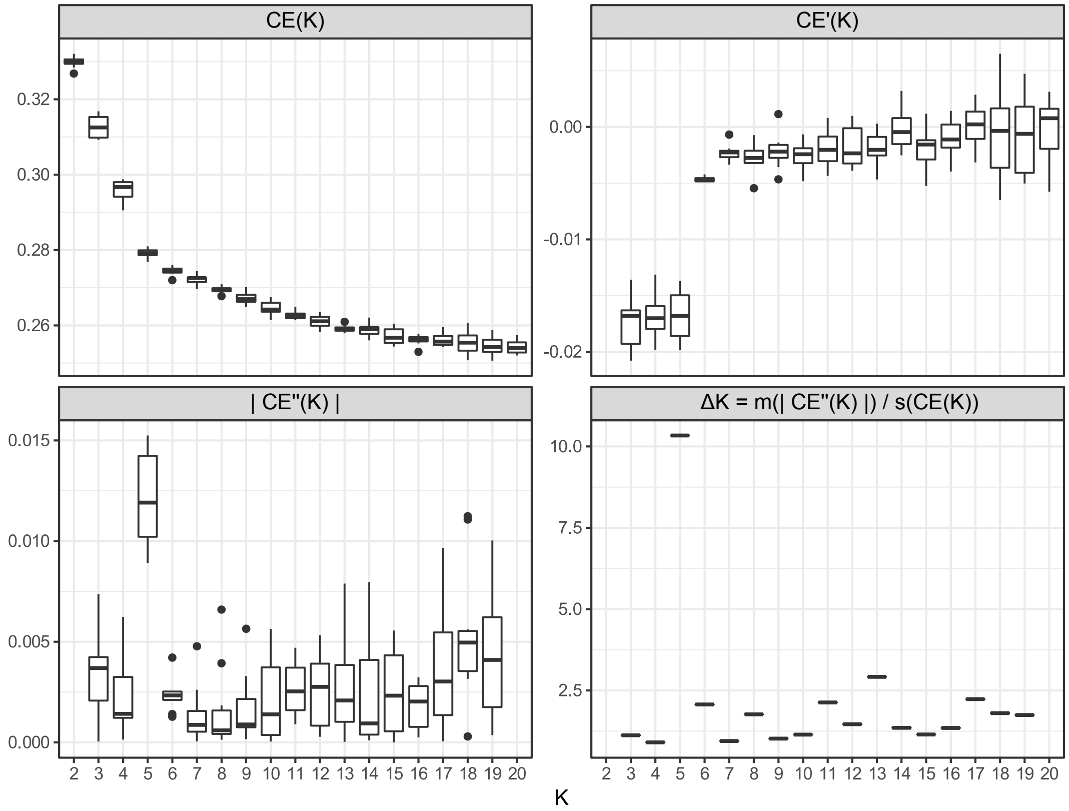

**Figure S4:** Rooted phylogeny with leaf labels and bootstrap values. This is the same tree as shown in Figure 2e of the article. The root is set to Cef2, a *B. stacei* accession collected in Sicily and sequenced for this study.

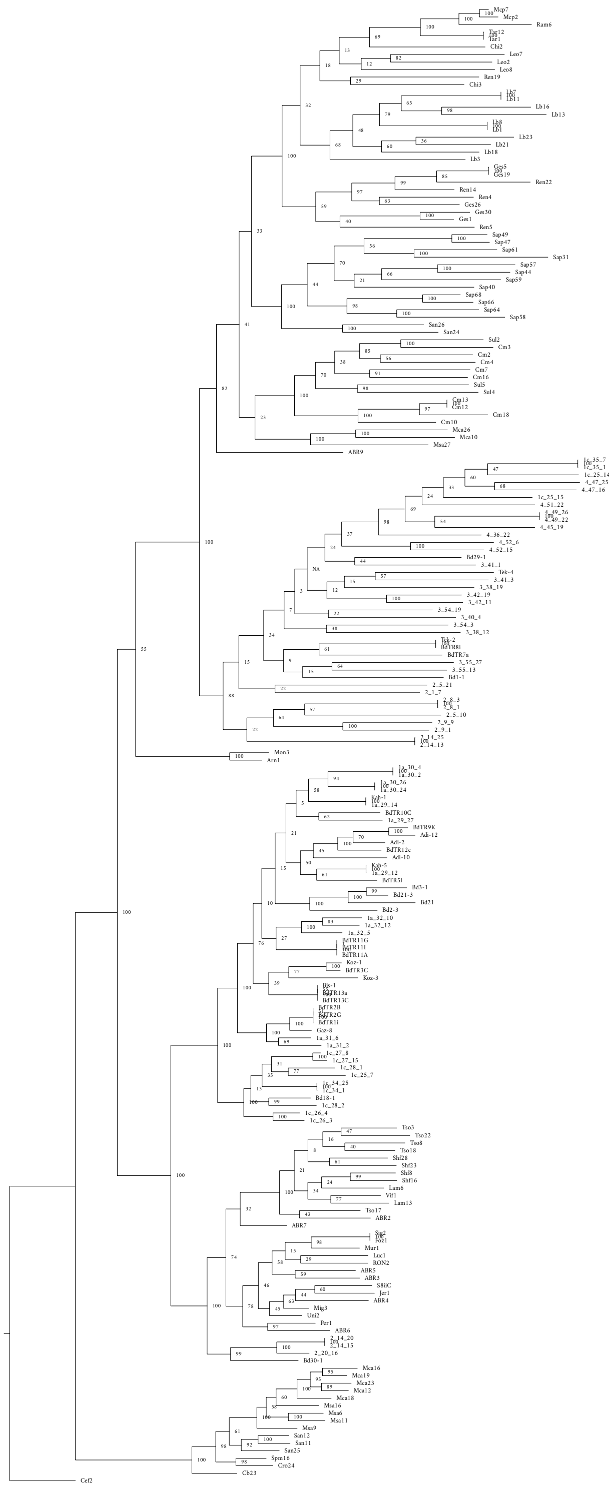

**Figure S5:** Sympatry of diverged lineages. Geographic details of the two locations where the A and the C lineage grow in sympatry. (a) Seven accessions sampled on Monte Calvo on the Gargano peninsula. (b) Five accessions sampled near Sanza in the Campania region.

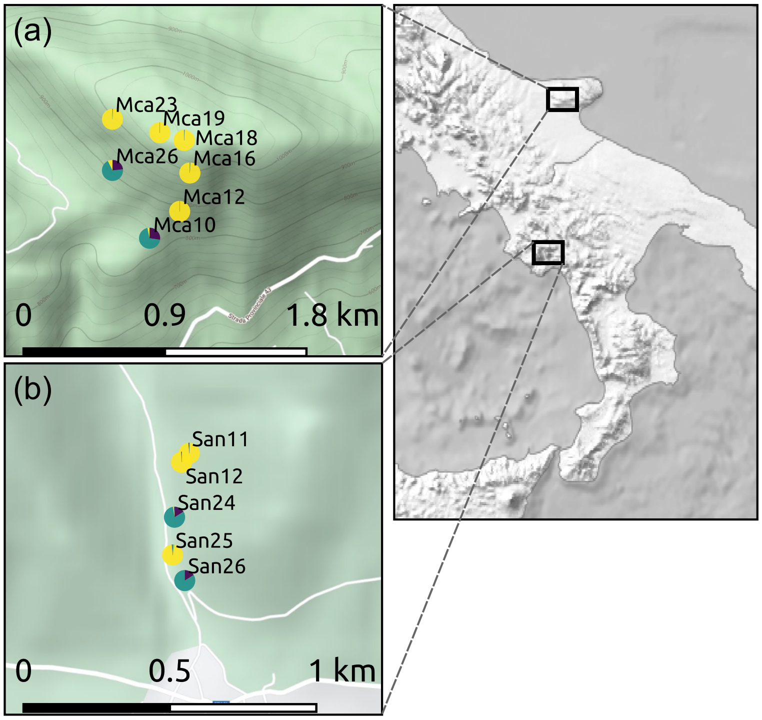

**Figure S6:** TreeMix analysis to test for gene flow between the five geographic clades. Panels on the left show the models and the model fit *f*, which represent the covariance of allele frequencies between populations explained by the model. Panels on the right display the residual fit.

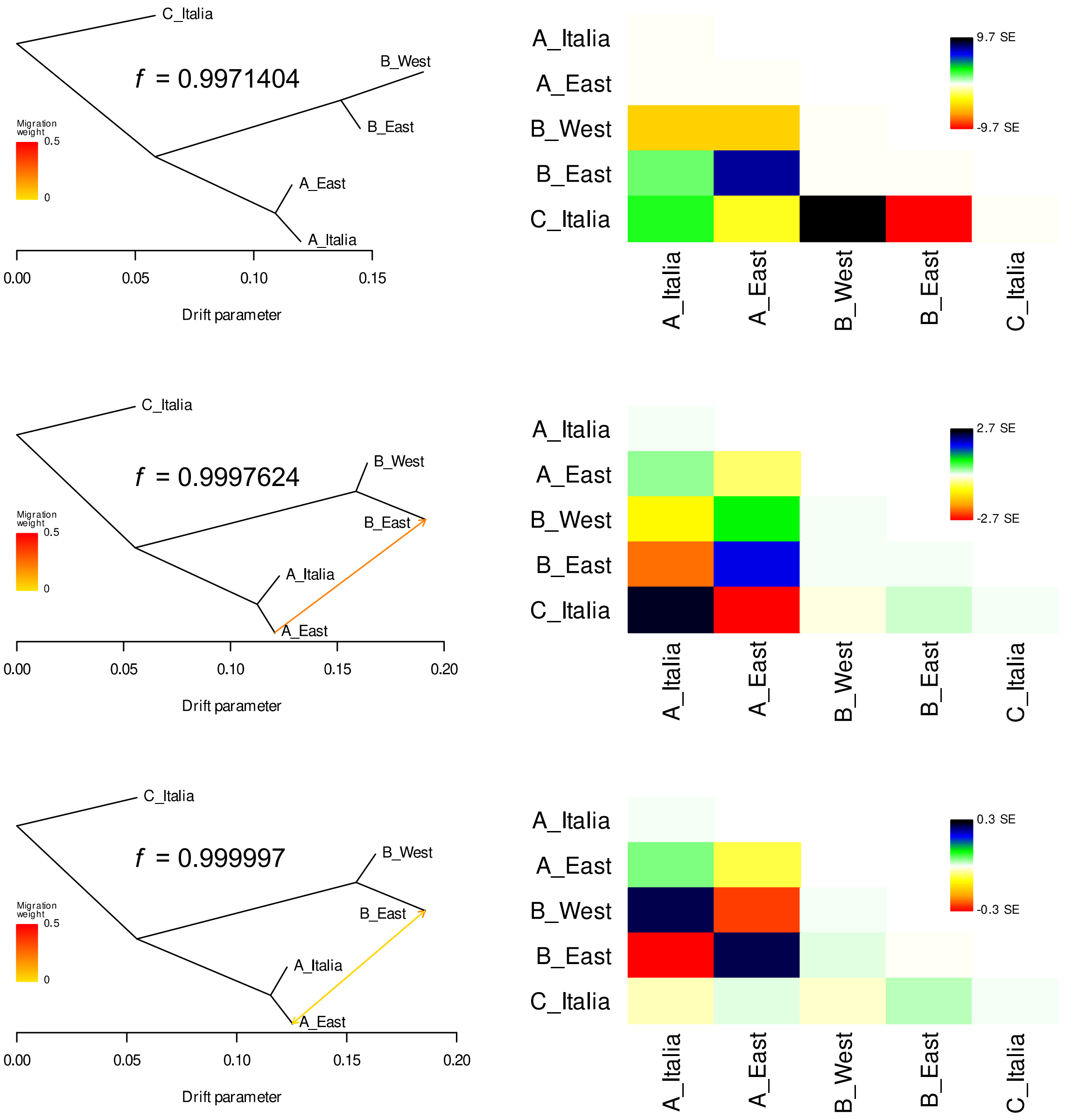

**Figure S7:** Relation between LD decay and terminal branch lengths in subtrees.

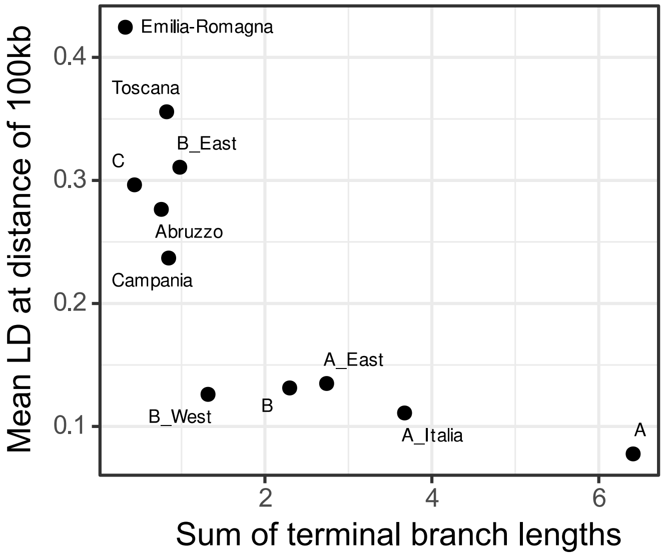

**Figure S8:** sNMF analysis with K from 2 to 20. Barplots are ordered according to the fineSTRUCTURE phylogeny, depicted on the left.

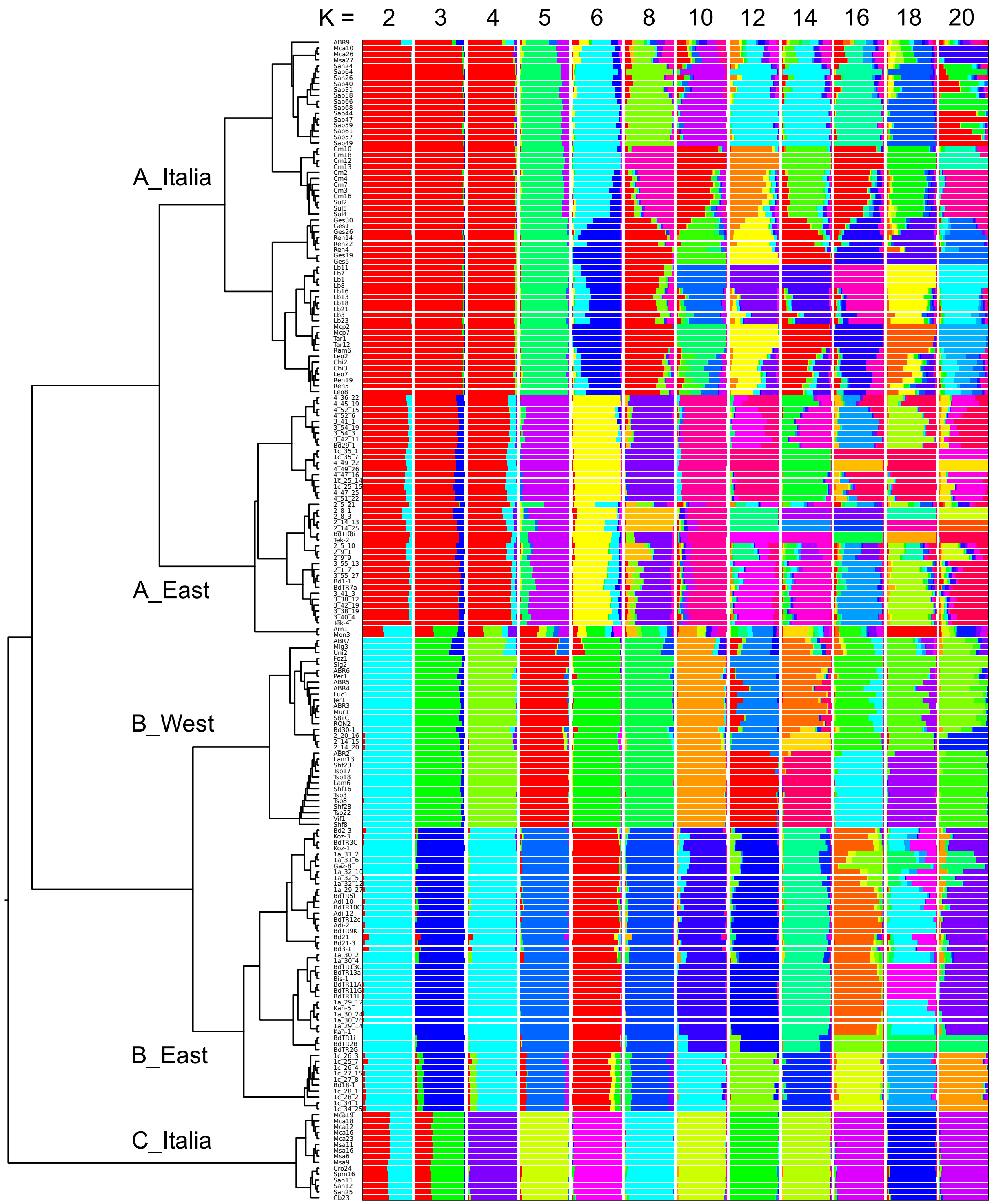

**Figure S9**: Within-clade PCAs. Seperate PCAs for each of the five geographic clades.

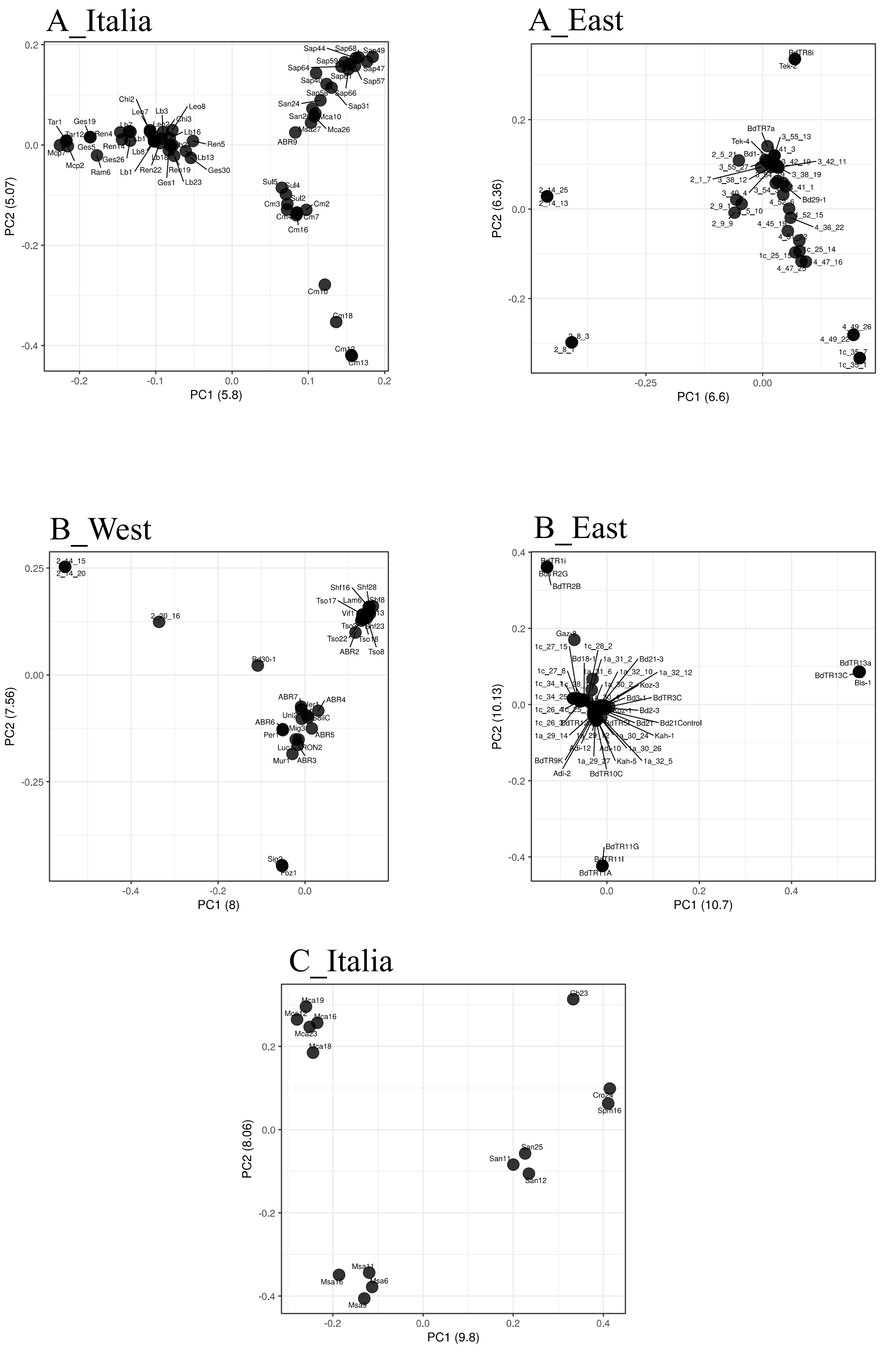

**Figure S10:** TreeMix analysis to test for gene flow between regional populations in Italy.

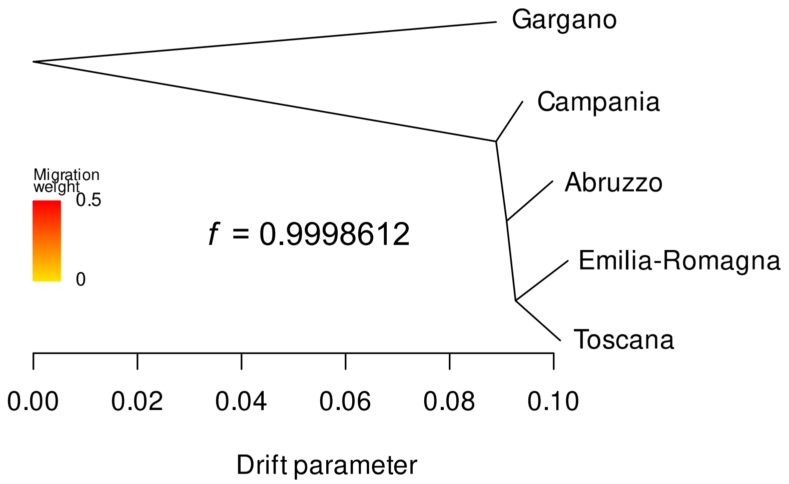


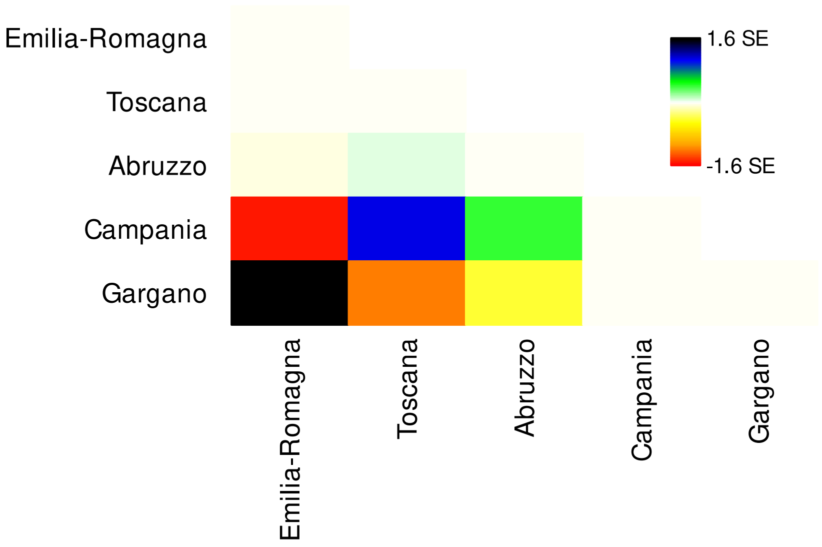

**Figure S11:** Flowering behaviour in the greenhouse (a) and under semi-natural conditions (b). These cumulative distributions show, for each lineage, the proportion of plants with flowers through time, counted in days since the appearance of the first flower in any lineage. a) Greenhouse experiment with 16 accessions and 3 replicates per accession, testing six different vernalization periods (0 to 12 weeks). No plants from the B_West lineage were tested, which in other experiments were shown to have an early flowering phenotype like the B_East accessions. b) Experiment under semi-natural conditions with 105 accessions and six replicates per accession. Tabulated results for the two flowering time experiments are available in Table S2.

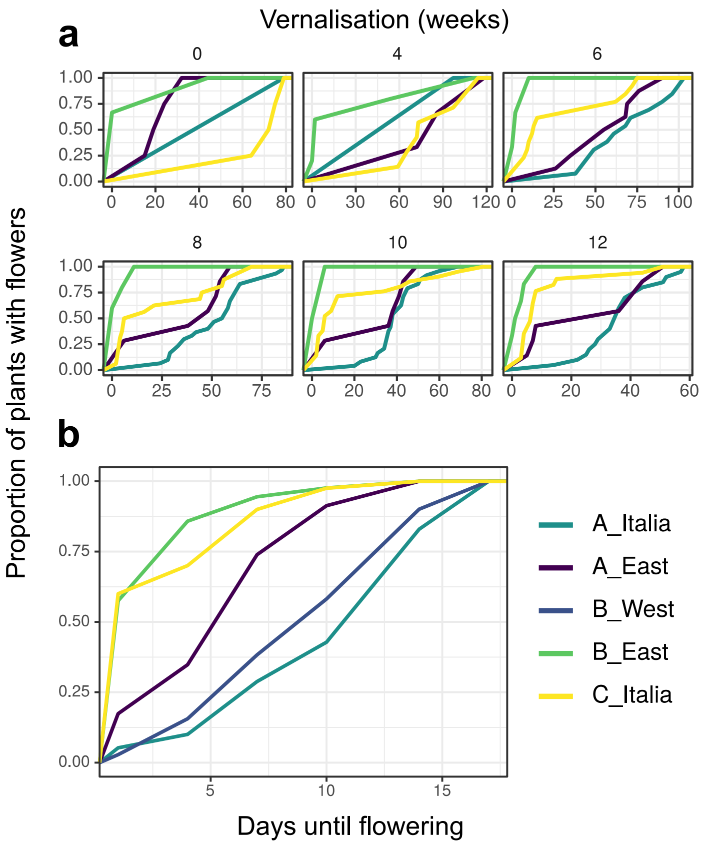
